# Fluctuating flooding motivates the facilitation of a wetland invader: a probe into the stress gradient hypothesis

**DOI:** 10.1101/2025.11.25.689875

**Authors:** Jingwen Zhang, Yang Qu, Xinran Li, Weiling Chen, Hongjin Xu, Jiyu Chen, Xue Yang, Yinuo Zhu, Tong Wang, Liyu Yang

**Affiliations:** Department of Forestry, College of Landscape Architecture and Forestry, Qingdao Agricultural University, Qingdao 266109, China; Shandong Peanut Research Institute/Chinese National Peanut Engineering Research Center, Shandong Academy of Agricultural Sciences, Qingdao 266100, China

**Author notes:** Corresponding author: *E-mail address:* (T. Wang).

**Keywords:** Wetland invaders · Plant interaction · Co-invasion · Invasion mechanism · Invasive species management

## Abstract

Stress gradient hypothesis proposes that the plant interaction tends to convert from competition to facilitation when facing accelerating stress. The present study aims to explore the interaction between two co-existing wetland plant invaders – *Alternanthera philoxeroides* and *Myriophyllum aquaticum* with exposure to flooding stress. Different plant assemblages including monospecific (single individual per species), homospecific (two individuals of same species) and heterospecific (two individuals of different species) treatments were subjected to different flooding regimes including mesic habitat, constant flooding and fluctuating flooding. The plant interaction was evaluated by biomass accumulation, biomass allocation and relative competition intensity index (RCI). We found that the biomass accumulation and allocation of *A. philoxeroides* were consistent across different treatments of community assemblage and flooding regime, while the root mass, biomass and root mass allocation of *M. aquaticum* significantly increased at the presence of *A. philoxeroides*. Moreover, the RCI of *M. aquaticum* decreased at the presence of *A. philoxeroides*. Especially, the RCI of *M. aquaticum* was 0.06 in the homospecific community, while that of *M. aquaticum* was -0.82 in the heterospecific treatment under fluctuating flooding condition. Overall, *Alternanthera philoxeroides* facilitated *M. aquaticum*, especially under fluctuating flooding condition, which complies with the stress gradient hypothesis. The water level drawdown can benefit the impedance of the growth *of M. aquaticum* and the deceleration of the co-invasion of *A. philoxeroides* and *M. aquaticum*.

## Introduction

The comparison between invasive and non-invasive counterparts has been explored extensively for detecting the invasion mechanism in the perspective of competition (Zhang et al. 2022; Gioria et al. 2023; Guo et al. 2023). Invasive species usually show a competitive edge such as faster growth than their non-invasive counterparts, which promotes their expansion in the introduced range (Kaushik et al. 2022; Li et al. 2024). As another facet of Darwin’s naturalization conundrum, invasive species can be similar with native species, which enables the invasive species to preadapt to the invaded habitats (Wang et al. 2023). However, the interaction of invasive species seems a forgotten piece of study on invasion ecology. From the point of competitive view, exotic species with superior competitiveness can exclude their exotic counterparts in the invasion process, which may cause an “invasional interference”, i.e. the reduction in the abundance or spread of co-existing exotic invaders (Rauschert and Shea 2017). However, the plant interaction not only includes the negative facet such as competition but also includes the positive facet such as facilitation (Brooker et al. 2008). The facilitation induces niche complementarity between plants and causes a win-win outcome (Brandt et al. 2023). Especially, a weaker competition is usually shown among co-existing exotic invaders than between the invaders and native species (Kuebbing and Nuñez 2016). Usually, the facilitation exists for species with niche differentiation, i.e. species with a niche overlap such as functional counterparts are usually difficult to form facilitation, since these species tend to compete with each other when they co-exist in the same habitat (Koffel et al. 2021; Pastore et al. 2021). For instance, Wang et al. (2019) reported that the co-invasion of two invasive free-floating plant species, *Pontederia crassipes* and *Pistia stratiotes* were realized through the resource niche differentiation in water nutrient use.

Environmental change can cause the reciprocal conversion between competition and facilitation. A widely-probed theory is that the plant interaction between co-existing individuals can shift from competition to facilitation with increasing abiotic stress (Callaway et al. 2002; Graff et al. 2007). In other words, the mutual benefit of different plants can be realized when the external environmental pressure intensifies, as postulated by the stress gradient hypothesis (Ali 2023). This theory is doubtful since limited resources likely inhibit the co-existence and promote the competitive exclusion. For instance, O’Brien et al. (2017) found that the severe rainfall deficit inhibited the species co-existence and caused the decrease of the plant productivity under nurse plant in a meta-analysis. Wang et al. (2022) reported that nitrogen addition and warming weakened the neighboring competitive effects on *Potentilla anserine* in assessing the interaction between two indicator species – *P. anserine* and *Ligularia virgaurea* of alpine meadows. However, as an explanation for the stress gradient hypothesis, the theory of niche expansion proposes that the nurse species can expand the resource niche for the beneficial species by facilitative approach such as secreting beneficial root exudates (e.g. metabolites relevant with plant growth promotion) and shaping beneficial soil microbiome (Rodrigues et al. 2021).

Flooding, as a unique disturbance type existing in wetlands, usually mediates stress for upland terrestrial plants as the stomata closure inhibits the CO_2_ diffusion into chloroplasts for photosynthesis (Martínez-Arias et al. 2022). However, the effects of flooding on wetland plants are complex considering the intensity and frequency of disturbance (Lan et al. 2021). Wetlands suffer more severe plant invasion than upland habitats and the clonal aquatic plants constitute a majority in the invasive plant category (Dudgeon et al. 2006). Prior studies commonly established a linkage between plant competition and the invasion success of exotic aquatic plant invaders (Huang et al. 2021; Hussner et al. 2021; Yuan et al. 2023). Many researches have revealed that the competitive advantage of exotic aquatic plant invaders over native plants (e.g. fast clonal growth and tolerance of harsh environments) is the key for their invasion success (Wang et al. 2018, 2023). The role of plant facilitation in determining the co-invasion success of wetland invaders has been verified in coastal wetlands (Zhang and Shao 2013). *Alternanthera philoxeroides* and *Myriophyllum aquaticum* are two common exotic freshwater plant species belonging to the same functional group, short clonal emergent plant. Based on our field investigation, these two species usually co-exist and form large mats. Therefore, we predict the existence of positive interaction between these two exotic aquatic plant invaders. Flooding, or the variation in hydrological regime is crucial in shaping the aquatic plant community, and also a key factor in determining the invasion success of exotic plant invaders (Wang et al. 2018). The present study subjected different community assemblages of *A. philoxeroides* and *M. aquaticum* to different flooding conditions and aimed to probe the intra- and inter-specific interaction between aquatic plant invaders in the face of environmental disturbance. The following predictions were postulated:

(1) A facilitation existed between the two exotic invaders;
(2) Flooding disturbance promotes the mutual benefit especially under fluctuating flooding condition, which corresponds with the stress gradient hypothesis.

## Materials and methods

### Plant materials

#### Alternanthera philoxeroides (Mart.) Griseb

*Alternanthera philoxeroides* (alligator weed), which originates from the Paraná River Basin of South America, currently have invaded the temperate, sub-tropical, and tropical China after approximately 90 years of invasion (Wang et al. 2018, 2023; Zhang et al. 2021). This species has terrestrial, semiaquatic, and aquatic growth-forms with great phenotypic plasticity however low genetic variance (Wang et al. 2005, 2018, 2023).

#### Myriophyllum aquaticum (Vell.) Verdc

*Myriophyllum aquaticum* (parrot’s feather) has been introduced from South America to East Asia for horticulture in 1920 and has invaded several European countries (Hussner and Champion 2012). Despite that *M. aquaticum* has not been officially affirmed to be invasive in China, the great invasive capacity has been considered by several publications (Wang et al. 2023; Zhang et al. 2021). This species has been observed to co-exist with *A. philoxeroides* in the field (Wang et al. 2018, 2023; Zhang et al. 2021). Water level is a crucial factor influencing its plant performance (Wang et al. 2018, 2023).

### Experimental set-up

On July 1^st^, 2024, 200 ramets for each of *A. philoxeroides* and *M. aquaticum* (approximately 30 cm in length) were collected from the riparian area of Moshui River of Qingdao City, China (120.40° E, 36.33° N) and were immediately transported to the experimental field of Qingdao Agricultural University (120.39° E, 36.32° N). Then, the 200 ramets of *A. philoxeroides* and *M. aquaticum* were respectively planted in 4 basins (diameter: 52 cm, height: 37 cm; 50 ramets per basin) with 10-cm-thick sediments collected from a lawn of the Chengyang Campus of Qingdao Agricultural University (ground, sieved and then evenly mixed). Tap water was added to each basin and a consistent water level of 1 cm was maintained. After 35 days of growth, we collected 72 ramets of each species (in total 144 ramets) with identical length of 15 cm from the basins on August 5^th^, 2024.

The collected 72 ramets were weighed for fresh weights and labeled. Then, the ramets were planted in 90 basins (diameter: 52 cm, height: 37 cm) with 10-cm-thick sediments collected from a lawn of the Chengyang Campus of Qingdao Agricultural University (ground, sieved and then evenly mixed). The experimental protocol was designed into five treatments as follows: 1) one individual of *A. philoxeroides* (monospecific), 2) one individual of *M. aquaticum* (monospecific), 3) two individuals of *A. philoxeroides* (homospecific), 4) two individuals of *M. aquaticum* (homospecific), and 5) one individual of *A. philoxeroides* and one individual of *M. aquaticum* (heterospecific). The water depth of each basin was consistent at 1 cm.

On October 1^st^, 2024, different treatments of flooding were operated after 55 days of growth. The flooding regime was designed as follows: 1) mesic habitat, 1 cm-deep water, 2) constant flooding, 25 cm-deep water, and 3) fluctuating flooding, the alternation of mesic and flooding conditions – 10 days of 25 cm-deep water, then 10 days of 1 cm-deep water, and then 10 days of 25 cm-deep water (Wang et al. 2018). Each treatment of planting design and flooding regime was repeated 6 times.

### Plant harvest

On October 30^th^, 2024, plant materials were harvested. For the treatments of mix culture including homospecific and heterospecific community assemblages, different plant individuals were separately harvested. After harvest, plant materials were separated into leaves, stolons and roots. The biomass of each part was determined after over-drying at 72 for 48 h. Then, the total biomass of each plant individual was calculated as the sum of leaf mass, stolon mass and root mass.

Biomass allocation including leaf mass ratio (LMR), stolon mass ratio (SMR), root mass ratio (RMR) and R/S ratio were calculated as follows:

LMR = Leaf mass / Total biomass

SMR = Stolon mass / Total biomass

RMR = Root mass / Total biomass

R/S ratio = Root mass / (Leaf mass + Stolon mass)

Competition intensity

The relative competition intensity index (RCI) was used to quantify the competition intensity of different species with different homospecific and heterospecific community assemblages (Grace 1995).

RCI=(P_mono_-P_mix_)/P_mono_

Where P_mono_ represents the biomass production of a plant in monoculture and P_mix_ represents the biomass production of a plant in mixture. When RCI>0, competition dominates; when RCI<0, facilitation dominates. The absolute value of RCI positively correlates with the intensities of competition or facilitation.

### Data analysis

All data of plant performance and biomass allocation met assumptions of normality and homogeneity of variance before analysis. A two-way ANOVA was used to analyze the effects of community assemblage and flooding regime on plant performance and biomass allocation of the two plant species. The Tukey’s test was performed to analyze the probable differences among different treatments. The above analyses were performed using SPSS 25.0 (IBM, Chicago, USA).

## Results

### Plant performance

The leaf mass, stolon mass, root mass and biomass of *A. philoxeroides* did not vary significantly across different treatments of community assemblage and flooding regime (Fig. 1a, b, c and d).

**Fig. 1.**
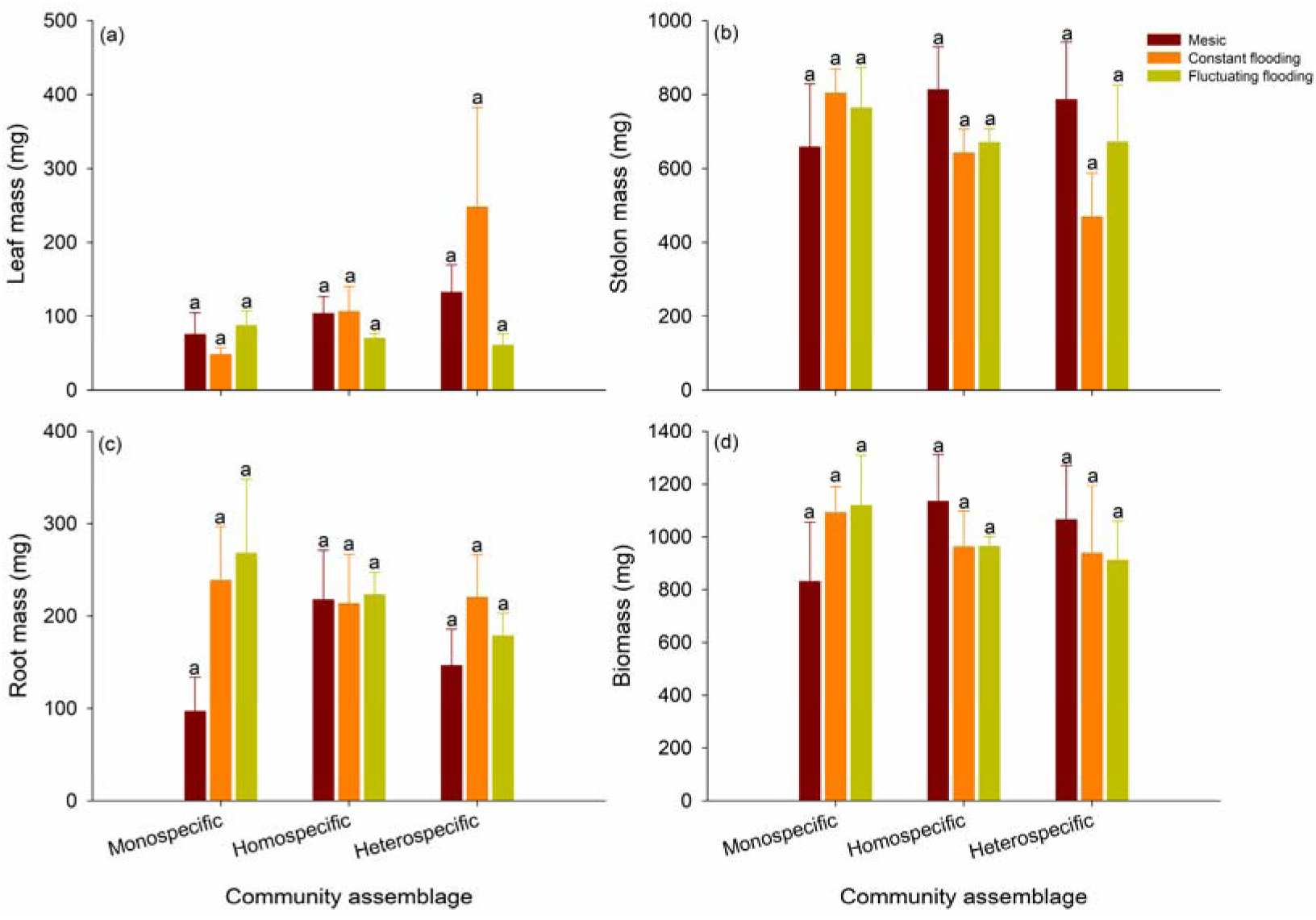
Plant performance of *Alternanthera philoxeroides* including: (a) leaf mass, (b) stolon mass, (c) root mass and (d) biomass in different treatments of community assemblage and flooding condition. Values are mean±SE. Bars with different lowercase letters represent significant difference at “*p*<0.05”

In the heterospecific community, the leaf mass of *M. aquaticum* was significantly greater in the fluctuating flooding treatment than in the constant flooding treatment (Fig. 2a). In the remaining treatments of community assemblage and flooding regime, the leaf mass, stolon mass, root mass and biomass of *M. aquaticum* did not vary significantly (Fig. 2a, b, c and d). However, the stolon mass, root mass and biomass of *M. aquaticum* were significantly greater in the heterospecific community than in the homospecific community (Fig. 2b, c and d). The root mass and biomass of *M. aquaticum* were significantly greater in the heterospecific community than in the monospecific treatment (Fig. 2c and d).

**Fig. 2.**
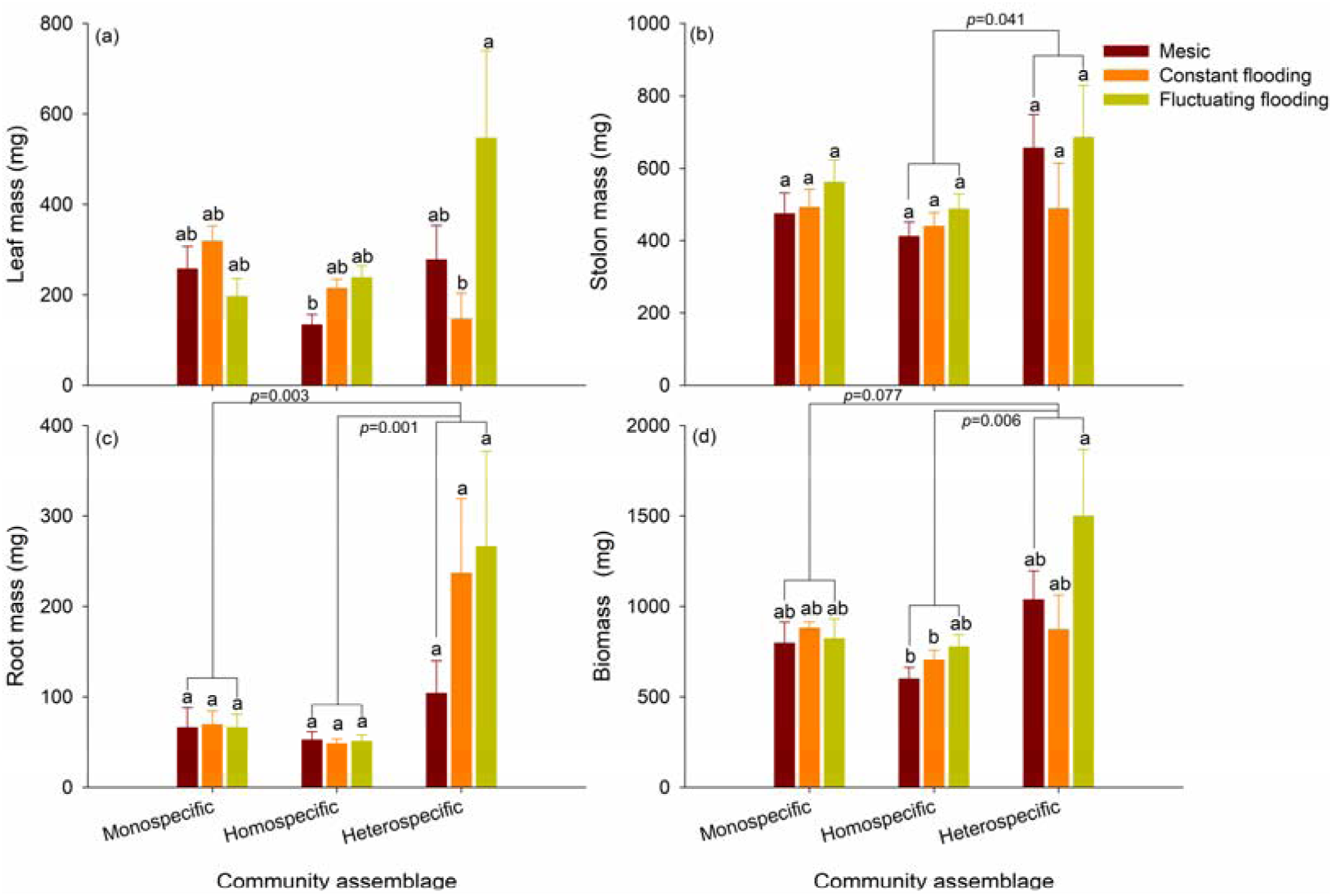
Plant performance of *Myriophyllum aquaticum* including: (a) leaf mass, (b) stolon mass, (c) root mass and (d) biomass in different treatments of community assemblage and flooding condition. Values are mean±SE. Bars with different lowercase letters represent significant difference at “*p*<0.05”

### Biomass allocation

The LMR, SMR, RMR and R/S ratio of *A. philoxeroides* did not vary significantly across different treatments of community assemblage and flooding regime (Fig. 3a, b, c and d). However, the SMR of *A. philoxeroides* was significantly greater in the monospecific treatment than in the heterospecific community (Fig. 3b).

**Fig. 3.**
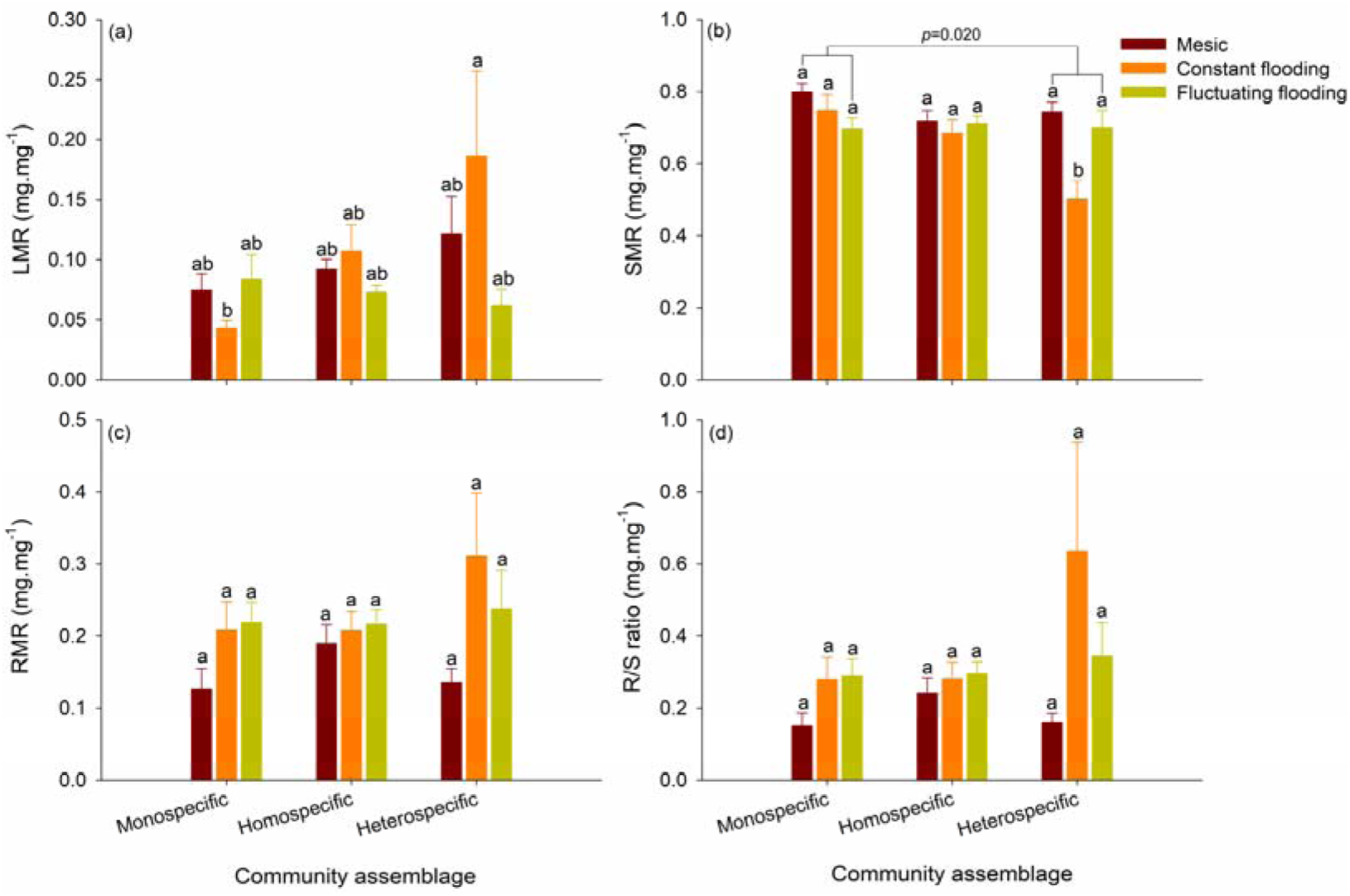
Biomass allocation of *Alternanthera philoxeroides* including: (a) LMR, (b) SMR, (c) RMR and (d) R/S ratio in different treatments of community assemblage and flooding condition. Values are mean±SE. Bars with different lowercase letters represent significant difference at “*p*<0.05”

The LMR and SMR of *M. aquaticum* did not vary significantly across different treatments of community assemblage and flooding regime (Fig. 4a and b). In the heterospecific community, the RMR and R/S ratio of *M. aquaticum* were significantly greater in the constant flooding treatment than in the mesic treatment (Fig. 4c and d). The RMR and R/S ratio of *M. aquaticum* were significantly greater in the treatment of fluctuating flooding treatment than in the remaining flooding treatments (Fig. 4c and d).

**Fig. 4.**
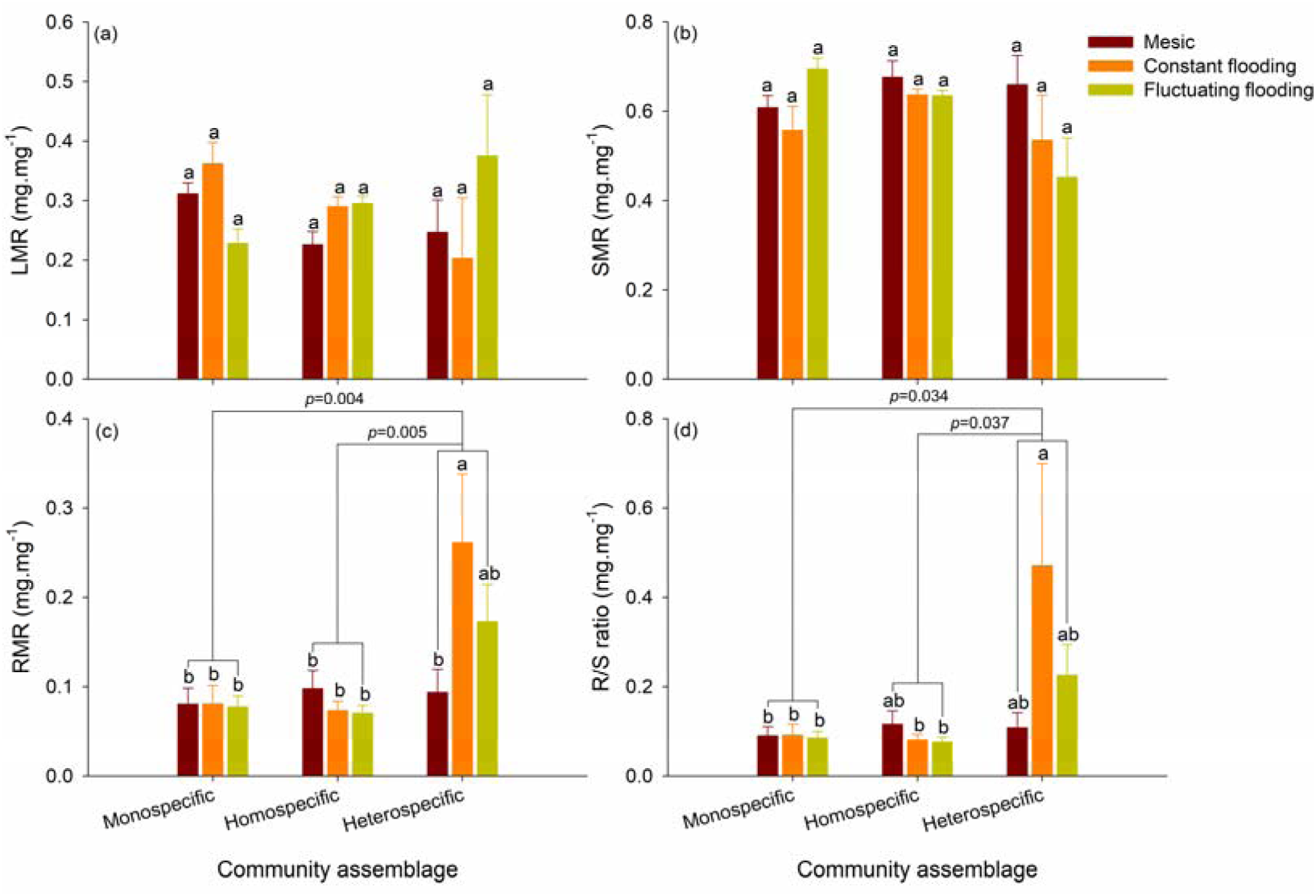
Biomass allocation of *Myriophyllum aquaticum* including: (a) LMR, (b) SMR, (c) RMR and (d) R/S ratio in different treatments of community assemblage and flooding condition. Bar with different lowercase letters represents significant difference at “*p*<0.05”

### RCI

The RCI values of *A. philoxeroides* were negative in the mesic treatment, while those were positive in the constant and fluctuating flooding treatments regardless of homospecific or heterospecific community assemblage (Fig. 5).

**Fig. 5.**
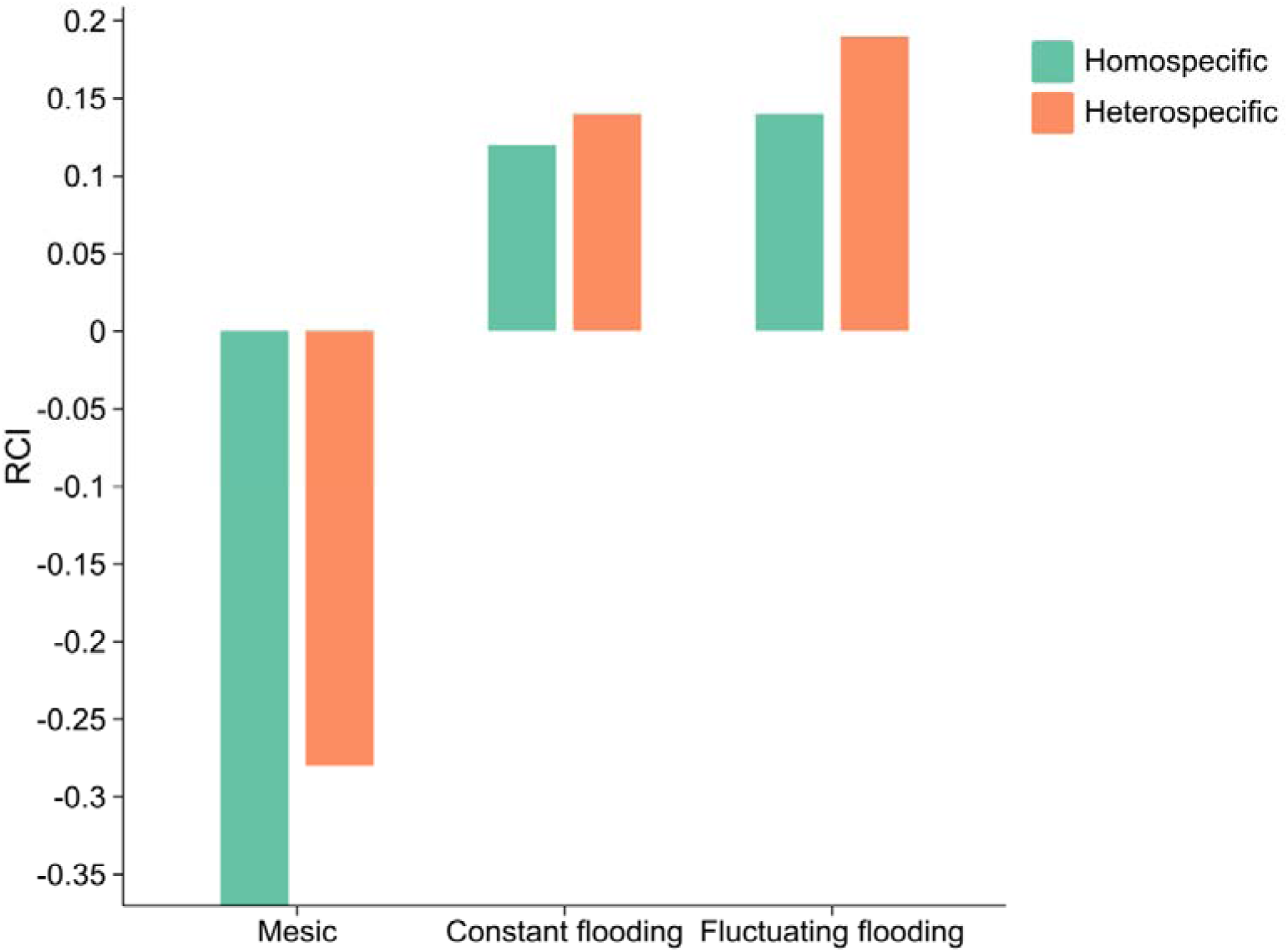
Relative competition intensity index (RCI) of *Alternanthera philoxeroides* in different treatments of community assemblage and flooding condition.

The RCI values of *M. aquaticum* were positive in the homospcific community across different flooding regimes, while those were negative in the heterospecific community across mesic and fluctuating flooding treatments (Fig. 6). Although the RCI value was positive in the heterospecific community and constant flooding treatment, the value was minimal at 0.01 (Fig. 6). Relative to the treatment of homospecific community, the treatment of heterospecific community led to the decreases of RCI value respectively at 0.55, 0.19, 0.88 (Fig. 6).

**Fig. 6.**
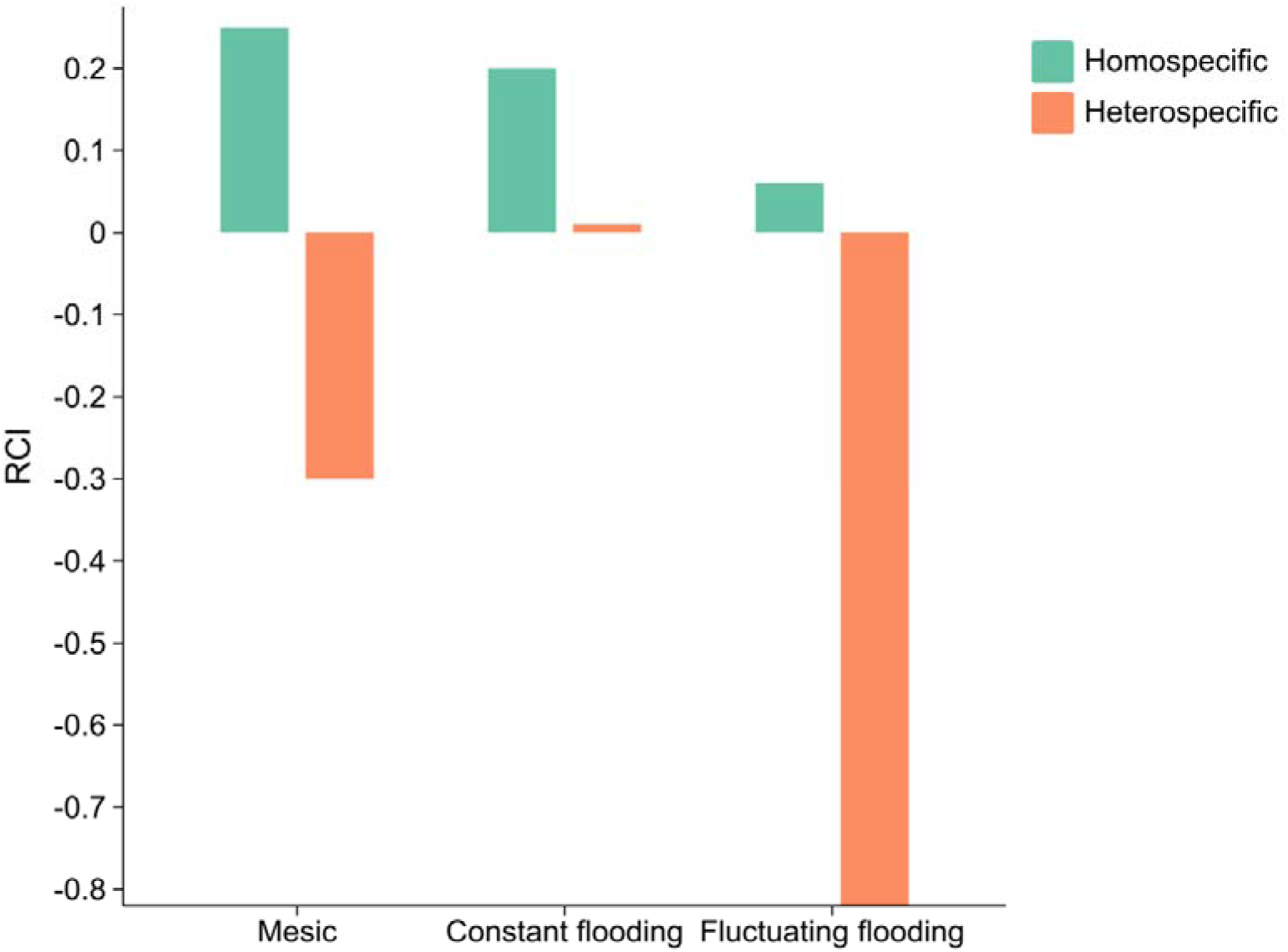
Relative competition intensity index (RCI) of *Myriophyllum aquaticum* in different treatments of community assemblage and flooding condition.

## Discussion

*Alternanthera philoxeroides* posed a facilitative effect on *M. aquaticum* Based on the niche theory of community ecology, an intense competition tends to exist for the functional counterparts (e.g. plants with similar life form or growth form) due to their niche overlap, i.e. a similar demand for resources (Koffel et al. 2021; Pastore et al. 2021). Following this paradigm, the intense competition between functional counterparts likely degrades the plant performance of any co-existing counterpart compared to monoculture. Moreover, con-specific individuals have same resource demands and experience a complete niche overlap, which should have experienced more intense competition than heterospecific individuals when co-existing in the same habitat (Barrero et al. 2024). However, the present study on the interaction of two invasive functional counterparts showed a contrast result as an invasive species benefited another invasive counterpart “costlessly”, which disobeyed these implications by the niche theory of community ecology. In the measurement of plant growth, we found that *A. philoxeroides*, the noxious wetland plant with a long invasive history (over 50 years) in China, enhanced the biomass accumulation of *M. aquaticum*, its functional counterpart expanding its invasive range in recent years. Nevertheless, a consistent plant performance of *A. philoxeroides* was shown among different treatments of community assemblage. Since root mass and root mass allocation significantly increased for *M. aquaticum* when co-existing with *A. philoxeroides*, the plant performance of *M. aquaticum* was facilitated owing to its root growth. Several explanations can be given: first, *Alternanthera philoxeroides* might secrete root exudates such as plant growth promoting metabolites that directly promote the root growth of *M. aquaticum*; second, *Alternanthera philoxeroides* probably modified the sediment microbiome through root physical intrusion and root exudates, which was beneficial for the root growth of *M. aquaticum* (Bongard 2012; Aizen and Torres 2024). In other words, *Alternanthera philoxeroides* promoted the growth of *M. aquaticum* via niche expansion (Rodrigues et al. 2021).

Considering RCI value, *A. philoxeroides* showed the same direction and similar magnitude in between the homospecific and heterospecific treatments, while the heterospecific treatment generally decreased the RCI values for *M. aquaticum* compared to the homospecific treatment. Therefore, the competitive intensity was similar for *A. philoxeroides* in different treatments of community assemblage; however, the competitive intensity showed an overall decreasing trend for *M. aquaticum* when co-existing with *A. philoxeroides*. Especially, in the treatments of mesic habitat and fluctuating flooding, the heterospecific treatment promoted the conversion of RCI values for *M. aquaticum* from positive (competition dominates) to negative (facilitation dominates) compared to the homospecific treatment (Callaway et al. 2002; Graff et al. 2007). Therefore, the evidence from RCI also verifies the beneficial effect of *A. philoxeroides* on *M. aquaticum.* In the field, the invasive *A. philoxeroides* hardly accompanies with other functional counterparts, and especially excludes the native wetland plants due to its strong clonal growth (Huang et al. 2025). Therefore, we usually observe that the invasive *A. philoxeroides* forms mono-stands or clonal mats via the asexual replication. However, the invasive *A. philoxeroides* can co-exist with other invasive wetland species for instance *M. aquaticum* in the field. The facilitative effect of *A. philoxeroides* on *M. aquaticum* partly revealed the reason *M. aquaticum* accompanying with *A. philoxeroides* (Soliveres et al. 2015).

Fluctuating flooding strengthened the facilitative effect of *A. philoxeroides* on *M. aquaticum* The stress gradient hypothesis hints two facets of plant interaction under disturbance: first, the disturbance facilitates the positive interaction between different species; second, the positive interaction between different species strengthens along with the increase of disturbance frequency and/or magnitude (Adams et al. 2022).

In the measurement of RCI, the present study partly supports the implications from the stress gradient hypothesis. First, fluctuating flooding promoted the decrease of RCI value in the heterospecific treatment for *M. aquaticum* relative to the remaining flooding treatments, suggesting the enhancing facilitative interaction under fluctuating flooding condition. However, the RCI value for *M. aquaticum* in the heterospecific treatment was positive under consistent flooding condition, while that was negative in the mesic habitat. Hence, the first implication from the stress gradient hypothesis was partly supported. We speculate that intermediate disturbance cannot exert prominent effects on the plant interaction between different species. Since intermediate disturbance may lead to the increase in available resources, the positive interaction may be not necessary for the maintenance of plant fitness (Jauni et al. 2015).

In the present study, although the consistent flooding caused the immersion of wetland plants and slowed the air diffusion in the water, the submergence can promote the wetland plants to recruit adventitious roots for water nutrient assimilation (Li et al. 2025; Pott and Pott 2021). Especially, *Myriophyllum aquaticum* is prone to form adventitious roots under submergence, which may weaken the facilitative effect of *A. philoxeroides* (Huang et al. 2024). However, the fluctuating flooding forces the wetland plants to switch growth forms (aquatic form with less clonal ramets and aquatic adventitious roots under submergence *vs.* wetland form with greater clonal growth and no aquatic adventitious roots) back and forth (Wang et al. 2018). The aid from facilitator becomes crucial for the fitness maintenance. In the present study, a negative RCI with the maximal absolute value was observed for *M. aquaticum* under fluctuating flooding. Considering the RCI distinction between homospecific and heterospecific treatments, the RCI value range was maximal in the treatment of fluctuating flooding (0.88 *vs.* 0.55 in the mesic habitat and 0.19 in under consistent flooding condition). Therefore, the observation from the RCI for *M. aquaticum* agreed with the second implication from the stress gradient hypothesis.

## Conclusions

The noxious invader, *A. philoxeroides* has been pervasive in China over its 90 years of invading history and has become a dominant wetland species excluding its native neighbors. Since *M. aquaticum*, despite an exotic species, has been increasingly used for waterscape architecture and wetland re-vegetation in China, its invasive potential may be augmented by the co-benefit from *A. philoxeroides* and global environmental change such as fluctuating flooding. Therefore, in order to maintain the ecological security of local wetland ecosystem, the introduction of *M. aquaticum* for ecological engineering especially with the existence of *A. philoxeroides* in the future should be prohibited. The artificial regulation of water level can be applied in the management of invasive wetland species. As the present study displays the intra- and inter-specific plant interaction of invasive wetland species, future studies should focus more on the mechanism beneath the observation. First, *A. philoxeroides* exerted positive effects on *M. aquaticum* without the impairment of performance itself. However, there should be a “cost” for itself. The multi-omics including transcriptomics and metabolomics may be helpful in digging out the “cost”. Second, as we postulated, *Alternanthera philoxeroides* may secrete root exudates to benefit *M. aquaticum* directly and indirectly.

Further explorations on root exudates and the role of them in altering the sediment microbiome are needed. Third, the role of aquatic adventitious roots in shaping the wetland plant interaction should be concerned, as the recruitment of adventitious roots under submergence modifies the mode for assimilating nutrients of wetland plants in the mesic habitats.

## Author contributions

Conducting the research, data analysis, data interpretation and writing were performed by Jingwen Zhang. Conducting the research was performed by Yang Qu, Xinran Li, Weiling Chen, Hongjin Xu, Jiyu Chen and Xue Yang. Conceptualisation, developing methods, data analysis, data interpretation and writing were performed by Tong Wang. Data analysis, data interpretation, preparation figures & tables and writing were performed by Yinuo Zhu and Liyu Yang. All authors read and approved the final manuscript.

## Conflict of interest statement

The authors declare that they have no known competing financial interests or personal relationships that could have appeared to influence the work reported in this paper.

**Table 1.** *F* values significances of two-way ANOVA of community assemblage and flooding regime on plant performance and biomass allocation of *Alternanthera philoxeroides* and *Myriophyllum aquaticum*.

|  | Community assemblage (C) |  | Flooding regime (F) |  | C × F |  |
| --- | --- | --- | --- | --- | --- | --- |
| *Alternanthera philoxeroides* | *F* | *p* | *F* | *p* | *F* | *p* |
| Leaf mass | 1.825 | 0.173 | 1.115 | 0.337 | 1.349 | 0.267 |
| Stolon mass | 0.549 | 0.582 | 0.693 | 0.505 | 1.060 | 0.388 |
| Root mass | 0.408 | 0.668 | 2.015 | 0.145 | 1.001 | 0.417 |
| Biomass | 0.069 | 0.934 | 0.005 | 0.995 | 0.675 | 0.613 |
| LMR | 2.905 | 0.065 | 1.429 | 0.250 | 2.095 | 0.097 |
| **SMR** | 5.784 | 0.006 | 6.829 | 0.003 | 4.148 | 0.006 |
| **RMR** | 0.803 | 0.454 | 4.080 | 0.024 | 0.990 | 0.423 |
| **R/S ratio** | 1.277 | 0.289 | 2.754 | 0.074 | 1.174 | 0.335 |
| *Myriophyllum*<br>*aquaticum* | *F* | *p* | *F* | *p* | *F* | *p* |
| **Leaf mass** | 2.082 | 0.137 | 1.773 | 0.182 | 3.200 | 0.021 |
| **Stolon mass** | 3.092 | 0.055 | 1.273 | 0.290 | 0.492 | 0.742 |
| **Root mass** | 9.208 | <0.001 | 1.082 | 0.348 | 1.111 | 0.363 |
| **Biomass** | 5.843 | 0.006 | 1.809 | 0.176 | 1.279 | 0.292 |
| **LMR** | 0.264 | 0.769 | 0.369 | 0.694 | 2.165 | 0.088 |
| **SMR** | 2.551 | 0.089 | 1.328 | 0.275 | 1.919 | 0.124 |
| **RMR** | 8.488 | 0.001 | 1.635 | 0.206 | 2.536 | 0.053 |
| **R/S ratio** | 4.743 | 0.014 | 1.470 | 0.241 | 1.846 | 0.137 |

## Acknowledgments

We sincerely appreciate the financial support from the Funding on Invasive Species Control from Shannxi Ganpo Ecological Technology & Engineering Co., Ltd (660/2424752) and the Open Project Funding of Key Laboratory of Intelligent Health Perception and Ecological Restoration of Rivers and Lakes, Ministry of Education, Hubei University of Technology (HGKFYBP07, 660/2424006).

## Data availability statement

Data are available from the authors upon reasonable request.

